# Enhancement of motor performance by a training method combining motor imagery and neurofeedback

**DOI:** 10.1101/2020.04.19.049783

**Authors:** Chosei Sha, Kaori Tamura, Tsuyoshi Okamoto

## Abstract

Motor imagery and neurofeedback have been proposed as motor training approaches, but their effects on the enhancement of motor performance are still controversial. This study aimed to enhance motor performance using a training protocol combining motor imagery and neurofeedback. Seventeen participants were randomly assigned to the training or control group. The training group received real-time electroencephalography activity feedback relative to the motor imagery of the motor action. The control group reiterated the motor imagery of the intended motor action without feedback. The motor performance of the training group was enhanced significantly more than that of the control group. Subsequently, the proposed training protocol was tested in five elite male tennis players to investigate its applicability to sports activities. The motor performance was enhanced in three of these five professional tennis players. Using our training protocol, which combined motor imagery and neurofeedback training, we achieved enhanced motor performance. Furthermore, we could suggest the applicability to sports from the results with world-level tennis players.

## Introduction

Motor skills are essential for daily life, and they need to be learned and developed to perform various activities. In sports, elite athletes must learn and develop specific motor skills at extremely high levels that are beyond those required for ordinary activities of daily living. To enhance motor performance, different training approaches have been investigated in many fields. In neuroscience, to enhance motor performance efficiently and effectively, motor imagery (MI) and neurofeedback (NFB) training have been proposed and studied.

The motor performance of a given action can be enhanced by the MI method, which consists of imagining the action without executing the movements involved (Decety and Grèzes 1999). In cases of rehabilitation, it has been reported that MI therapy supplements conventional physiotherapy or occupational therapy for patients with motor dysfunction disorders (Caligiore et al. 2017; Zimmermann-Schlatter et al. 2008). These studies have reported improvement of the movements of the upper and lower limbs among patients following stroke. Conversely, in studies involving healthy participants, MI training can yield effects that enhance motor performance (Gentili et al. 2006; Ruffino et al. 2017, 2019; Sobierajewicz et al. 2017). These studies have reported that the duration of an actual movement is decreased by MI training. The above studies among patients and healthy participants suggest that MI has the potential not only to improve the patients’ motor functions but also to enhance the motor performance of healthy individuals.

Among the many investigations of MI training, researchers noted that its effect seems to be variable. A review analyzed 133 MI training studies with positive, unchanged, or negative results (Schuster et al. 2011). The analysis suggested that successful MI training could be achieved with an optimal protocol. However, the authors lacked a neurophysiological viewpoint. Di Rienzo *et al.* have discussed the effects of MI training on activity-dependent neuroplasticity (Di Rienzo et al. 2016). We agree that MI training protocol needs to be optimized, but this end would be better served if we first understand how MI can modulate neurophysiological activities.

The characteristic brain activity of MI is an event-related desynchronization (ERD) of mu-band (8–13 Hz) oscillation in sensory-motor areas. This mu-band ERD has also been observed during actual movement execution(Sobierajewicz et al. 2017). Many studies have shown that mu-band ERD was induced in the contralateral hemisphere of the hand during MI (Wolf et al. 2014; Wriessnegger et al. 2018). In sports, some studies have reported that mu-band ERD of elite athletes was larger than that of amateurs or novices during the MI of sports activity (Del Percio et al. 2007; Wolf et al. 2014). The differences of mu-ERD between elite and non-elite athletes have shown some correlation but not a causal relationship.

In most MI studies, the mu-ERD has been used to monitor the mental states of subjects during MI (Kim et al. 2017; Wriessnegger et al. 2018). It has been reported that the vividness of MI correlates with mu-ERD amplitudes (Toriyama et al. 2018). The feedback of mu-ERD will enable the trainees to confirm the vividness of MI by themselves, which will lead to an enhancement of motor performance (Ruffino et al. 2017). For effective training, the vividness of MI estimated by mu-ERD should be conveyed to a trainee rather than a trainer.

To regulate motor-related neuronal activity, we overlaid NFB training on MI training. NFB training is a method of self-regulation of neural activities to control specific mind or brain states. In the NFB training paradigm, brain activities are transformed into external sensory stimuli, which are fed back continuously. Individuals can learn to regulate their brain states by receiving this continuous feedback. Several studies have reported the enhancement or improvement of cognitive function (Hanslmayr et al. 2005; Kober et al. 2019; Nan et al. 2012) with feedback signals of sensory-motor rhythm (SMR, 12–15 Hz) (Kober et al. 2019) and alpha band (individual alpha frequency [Nan et al. 2012] or common frequency of 10–13 Hz [Hanslmayr et al. 2005]).

Although NFB training using SMR (13–15 Hz) and alpha-theta: 4–8 Hz (Rostami et al. 2012) has been used in many types of sports, the effects vary across sports and individuals (reviewed in Gruzelier 2014; Jeunet et al. 2019). In an NFB-based study on archery, focusing on brain activities peculiar to elite athletes, pre-elite athletes were trained using the feedback of the EEG activity in the left temporal site (in which the elite athletes showed higher activation during shooting) and had a higher enhancement of shooting accuracy in the NFB group than in the sham or no feedback groups (Park et al. 2015; Thompson et al. 2008). An NFB study of table tennis players evaluated SMR (12–15 Hz) in 10 elite athletes and reported that some athletes showed an increase in accuracy that was not observed in other athletes (Brown et al. 2012). Further, in an NFB study of golf players a personalized EEG profile was determined during successful golf putts during a 3-day NFB training using individually assessed measures. The results showed that their performance was significantly enhanced from day 1 to day 2; however, they could not find significant enhancement on day 3 (Arns et al. 2007). To ensure stable effects, an effective training protocol should be developed.

The large variation in the effects on motor performance may be caused by two factors: the training target (the kind of brain activity occurring during training) and the training strategy (the strategy used to change the neuronal activity during NFB training). In previous NFB studies on sports, training targets of EEG activities were not directly related to the intended motor action, and training strategies were not standardized by experimenters and were possibly unrelated to the intended action because strategies were selected by the participants. In most NFB studies, including in those relevant to sports training, the strategies have been freely determined by the trainee (Paret et al. 2019). To enhance motor performance efficiently, the training target and training strategy should be directly associated with the intended motor action.

Our main purpose is to determine whether the mu-band ERD enhancement can improve motor performance. If the mu-band ERD differences between athletes and novices are the result of actual motor training (Del Percio et al. 2007; Wolf et al. 2014), training of mu-band activity will not provide any enhancement of motor performance. If motor performance and mu-band activity can affect each other, motor performance will be enhanced by mu-band training. In this study, we used the term “mu-ERD” to define the mu-band activity induced by MI. Here, we hypothesized that mu-ERD feedback during MI will regulate motor-related ERDs, and the enhancement can improve actual motor performance. For mu-ERD training, real time feedback of motor-ERD during MI will be effective using NFB methods.

In addition to mu-ERD, beta-ERD is another MI-related ERD (Paret et al. 2019). Many studies have proposed that beta-ERD is also related with motor planning, execution, and imagery (Formaggio et al. 2010; Kraeutner et al. 2014; Nasseroleslami et al. 2014; Neuper et al. 2006; Gert Pfurtscheller and Neuper 1997; Wheaton et al. 2009). Some reports have shown a correlation of beta-ERD with motor performance (Gehringer et al. 2019; Jochumsen et al. 2017) and motor learning (Pollok et al. 2014), but others have shown a negative correlation between beta-ERD and performance (Dal Maso et al. 2018; Gehringer et al. 2018). To limit the effect of neurofeedback, we focused on the mu-ERD feedback and not beta-ERD. We performed an offline analysis of the transfer effect on beta-ERD enhancement in the proposed protocol.

Based on our hypothesis that motor-related ERD enhancement can enhance motor performance, we developed a training protocol combining MI and NFB (MI-NFB). This protocol was designed to enable the training of vividness of motor imagery and mu-ERD regulation at the same time, and to provide more stable effects than either MI or NFB alone. When trainees imagined a given motor action under the protocol, they received mu-ERD feedback related to the action. The training strategy consisted of imagining the intended motor action to enhance its performance.

MI-NFB approach has been demonstrated in patients and healthy persons (*e.g.*, Spychala et al. 2020; Wriessnegger et al. 2018; Zich et al. 2017, reviewed in Jeunet et al. 2019). In the previous studies of patients, their purpose was to restore the patients’ motor functions to normal or near-normal levels, and the target movements were designed for rehabilitation (Spychala et al. 2020; Zich et al. 2017). In studies of healthy persons, the MI-NFB approach has been investigated with several neuroimaging techniques such as Electroencephalogram (EEG) (Bai et al. 2014; Wriessnegger et al. 2018; Zich, De Vos, et al. 2015; Zich, Debener, et al. 2015), Magnetoencephalogram (MEG) (Boe et al. 2014), functional Magnetic Resonance Imaging (fMRI) (Zich, Debener, et al. 2015), and Near-infrared spectroscopy (NIRS) (Kober et al. 2014, 2018). Furthermore, some reports have examined frequency band changes associated with MI (Kraeutner et al. 2014; Wriessnegger et al. 2018).

It is worth noting that most MI-NFB studies have aimed to develop a brain computer interface or to optimize MI parameters. Fewer investigations have aimed to evaluate the enhancement of motor performance through training. The aims of this study were, first, to enhance motor performance by MI-NFB training and, second, to enhance MI ability or mu-ERD. Additionally, the proposed protocol was carried out in a limited number of elite athletes to test the possibility of its application and to examine each case.

Our objective was to demonstrate that our novel MI-NFB training protocol can enhance motor performance more efficiently than conventional MI training. First, we verified the effectiveness of the training protocol in a laboratory with university students. The unfamiliar motor action was designed originally as a pegboard game to evaluate the effect of MI-NFB on motor performance itself superior to MI ability or motor-related ERDs. Subsequently, we investigated its applicability to sports performance with a small sample of world-class tennis players as a preliminary study. The tennis serve was the motor action enhanced by the training protocol.

## RESULTS

### Experiment 1: Training protocol for healthy volunteers

Participants were assigned to two groups: the MI-NFB group and the control group. The motor performance and related neural activities were analyzed according to the performance of the dominant and non-dominant hand separately. Before starting the training, their general MI ability was assessed using the Kinesthetic and Visual Imagery Questionnaire (KVIQ) (Kober et al. 2018). Visual imaging score (VIS), kinesthetic imaging score (KIS), and the sum of KIS and VIS (SS) were analyzed (Malouin et al. 2007). There was no significant difference between the groups in VIS (t = −0.71, *P* = 0.49), KIS (t = −1.0, *P* = 0.34), and SS (t = −0.89, *P* = 0.39) before the training.

### Clustering analysis

Many previous studies have reported that motor performance, even in skilled motor activities, is impaired by mental fatigue (Marcora et al. 2009; Schiphof-Godart et al. 2018). To verify the effects of the proposed training protocol without the influence of the harmful effects caused by mental fatigue, clustering analysis was performed for the fatigue reports and motor performance data. We used a data-driven method (a hierarchical cluster analysis of the mental fatigue during MI training in all participants) (see Fig. 1A and 1B for the detailed results). As a result, four participants who did not show a decrease in fatigue levels in the 3-day period were classified into another cluster (no-habituation group) (Fig. 1C), and they all underwent MI-NFB training. No one who performed a control task was classified as a no-habituation participant. The remaining data were separated according to the performed task into MI-NFB and control.

**Fig. 1.**
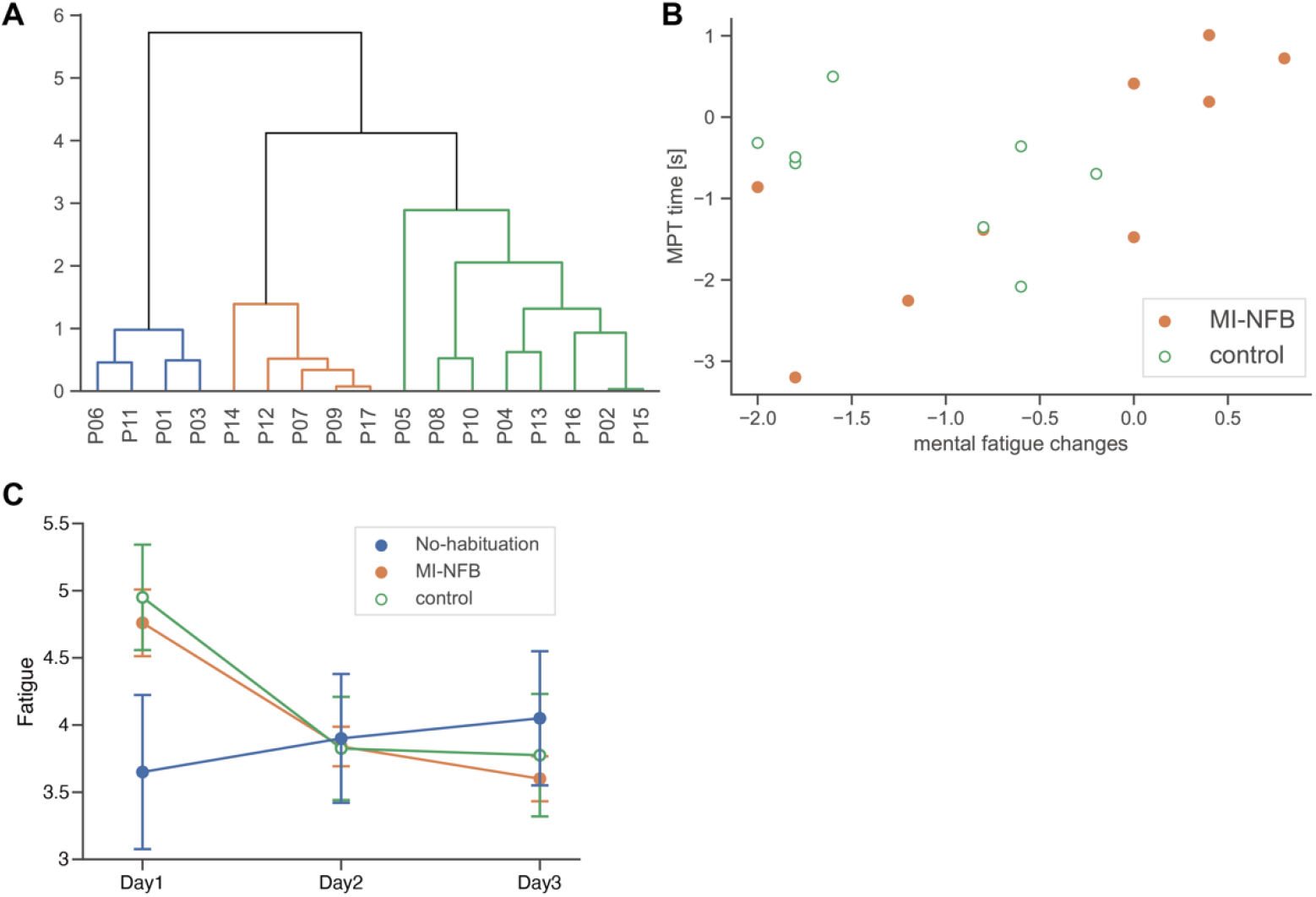
A: Dendrogram of the hierarchical clustering analysis of mental fatigue and motor performance test (MPT) working time. B: Scatter plots showing changes in mental fatigue according to MPT working time. C: Mean and SEM of mental fatigue in each group

### Dominant hand

We compared the working time of the motor performance test (MPT) using a pegboard before and after the training and analyzed by repeated measures ANOVA (pre-/post-training × groups). There was a significant main interaction effect (F(2, 97) = 10.9, *P* = 0.0017). The multiple comparison showed a significant time-series change in the MI-NFB group (F(5,4) = 3.87, *P* = 0.013, α = 0.017), but not in the control or no-habituation groups (F < 2.24, *P* > 0.10). The multiple comparison showed a significant time-series change in the MI-NFB group (t = 4.51, *P* = 0.011, α = 0.017), but not in the control or no-habituation groups (*P* > 0.047, α = 0.017) (Fig. 2A).

**Fig. 2.**
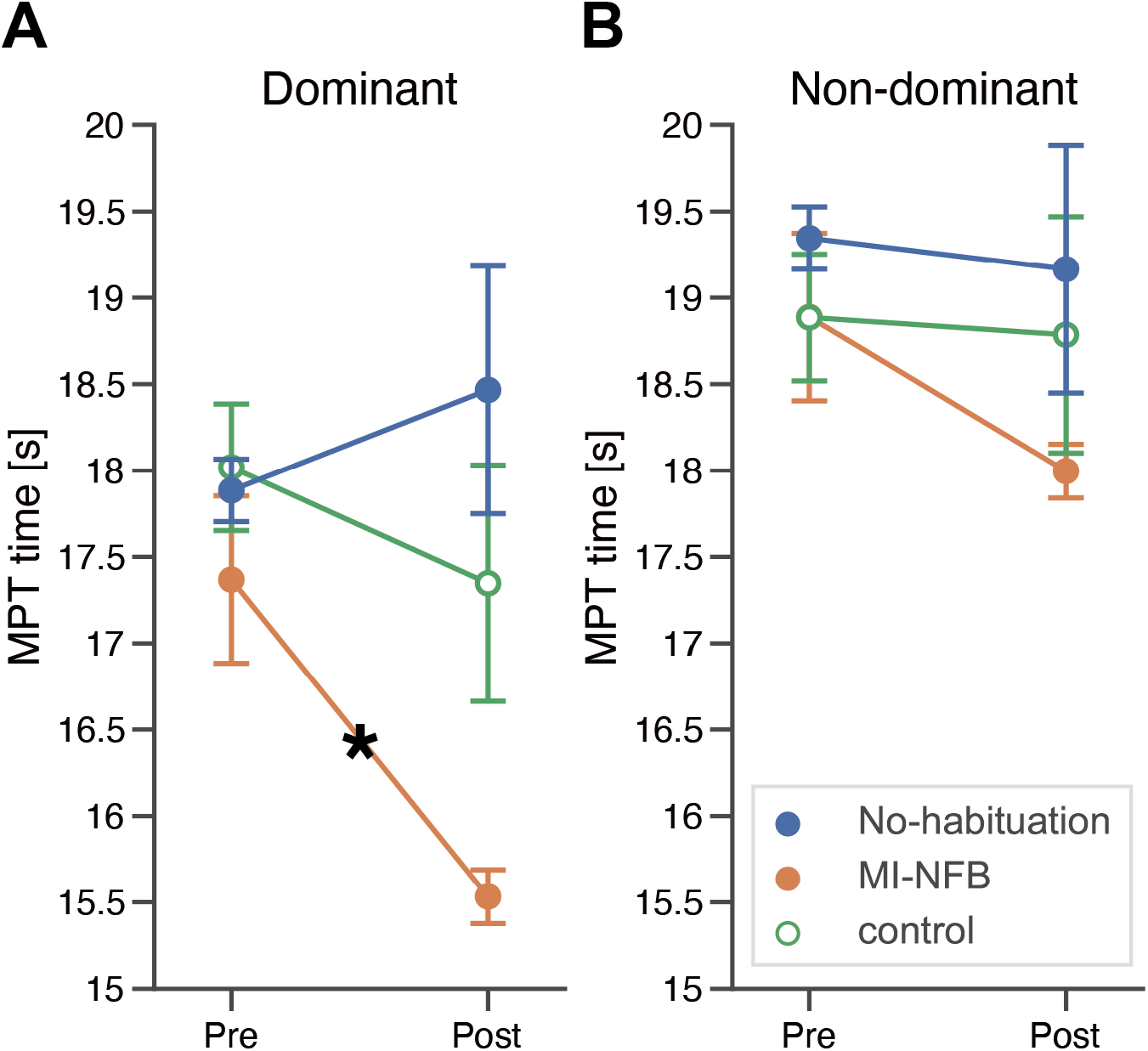
Mean and standard error of mean (SEM) of the working time of the motor performance test (MPT) with the dominant (A) and non-dominant hands (B) between pre and post training. *: *P* < 0.017, with a Bonferroni correction. MI-NFB: mental imagery-neurofeedback

To evaluate the training effect, we analyzed the changes of mu-ERD (8–13 Hz) during the evaluation sessions using a repeated measures ANOVA (pre/post on the three training days × groups). There was no significant main effect of groups (F(2, 13) = 2.78, *P* = 0.10), but a marginal main effect of the repetitions was observed (F(5, 78), *P* = 0.041). There was no significant interaction effect (F(10, 65) = 1.35, *P* = 0.22). (Fig. 3A). The mu-ERD distributions are shown in Fig. 3B.

**Fig. 3.**
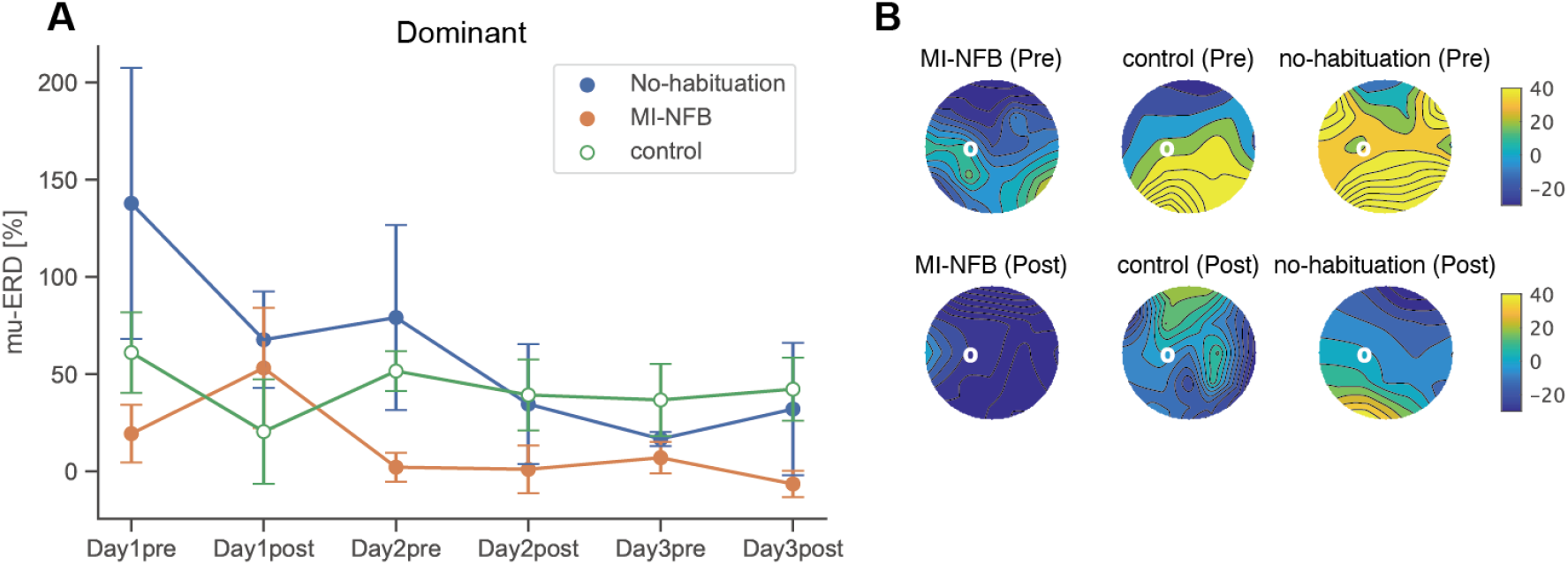
Results of mu-ERD with the dominant hand A: Mean and SEM in the mu-ERD from the pre-training session on day 1 to post-training on day 3. B: Topography of the grand averaged mu-ERD in each group. Color bars show ERD [%]. White circles indicate channel location of C3, which was used as feedback electrode. ERD: event-related desynchronization, MI-NFB: mental imagery-neurofeedback

To investigate the transfer effect on beta band, we analyzed the beta-ERD changes (13–28 Hz). A repeated measures ANOVA (pre/post on the three training days × groups) showed a significant main effect of group (F(2,13) = 6.27, *P* = 0.012) and a marginal significant interaction effect (F(10,65) = 1.87, *P* = 0.086). The multiple comparison showed a significant time-series change in the MI-NFB group (F(5,4) = 3.87, *P* = 0.013, α = 0.017) but not in the control or no-habituation groups (F < 2.24, *P* > 0.10). Post-hoc test showed a significant difference of beta-ERD between pre-training on day 1 and post-training on day 3 in the MI-NFB group (t = 4.26, *P* = 0.013) (Fig. 4A). The beta-ERD distributions were indicated in Fig. 4B.

**Fig. 4.**
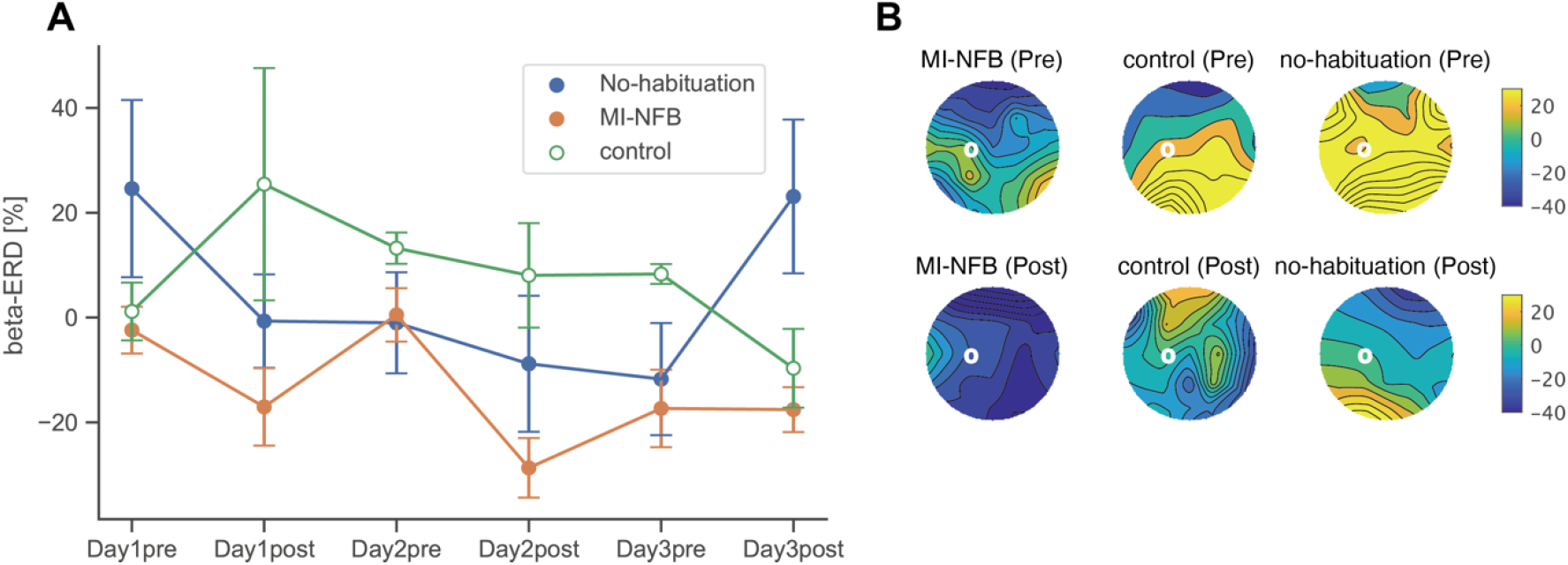
Results of beta-ERD with the dominant hand A: Mean and SEM in the mu-ERD from the pre-training session on day 1 to post-training on day 3. B: Topography of the grand averaged mu-ERD in each group. Color bars show ERD [%]. White circles indicate channel location of C3, which was used as feedback electrode. ERD: event-related desynchronization, MI-NFB: mental imagery-neurofeedback

### Non-dominant hand

The repeated measures ANOVA showed no significant main effect or interaction (pre-/post-training × groups) in the working time of MPT using the non-dominant hand (F < 1.10, *P* > 0.31) (Fig. 2B).

To evaluate the training effect, we compared the mu-ERD during the evaluation session using a repeated measures ANOVA (pre/post on the three training days × groups). There was no significant main effect of group (F(2, 13) = 0.37, *P* = 0.70) and no significant interaction effect (F(10, 65) = 0.68, *P* = 0.74) (Fig. 5).

**Fig. 5.**
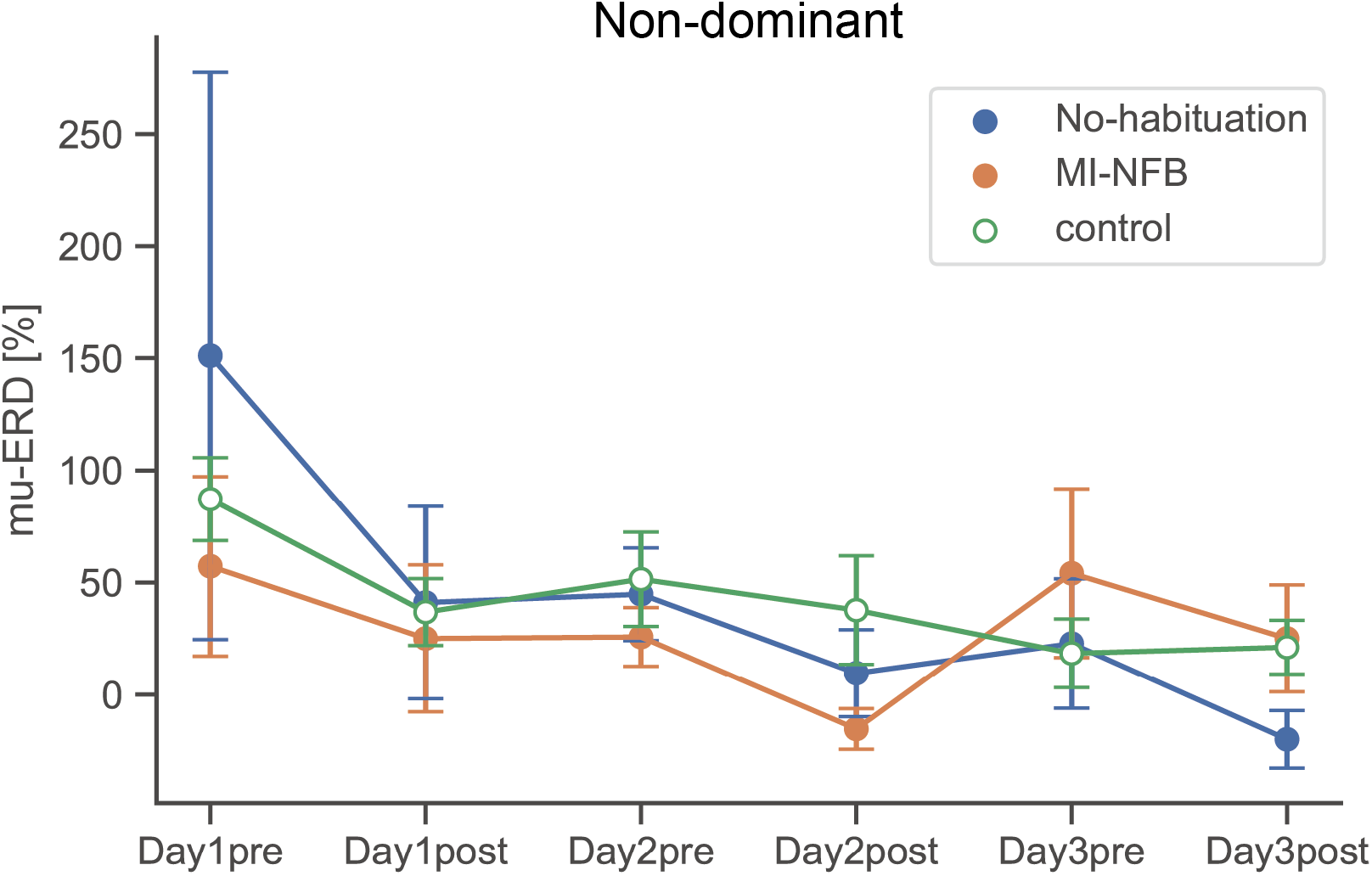
Results of mu-ERD with the non-dominant hand Mean and SEM in the mu-ERD from pre-training on day 1 to post-training on day 3. ERD: event-related desynchronization, MI-NFB: mental imagery-neurofeedback

The beta-ERD changes were analyzed in the dominant hand. A repeated measures ANOVA (pre/post on the three training days × groups) showed no significant main effect of group (F(2, 13) = 2.48, *P* = 0.12). There was no significant interaction effect (F(10, 65) = 0.88, *P* = 0.56) (Fig. 6).

**Fig. 6.**
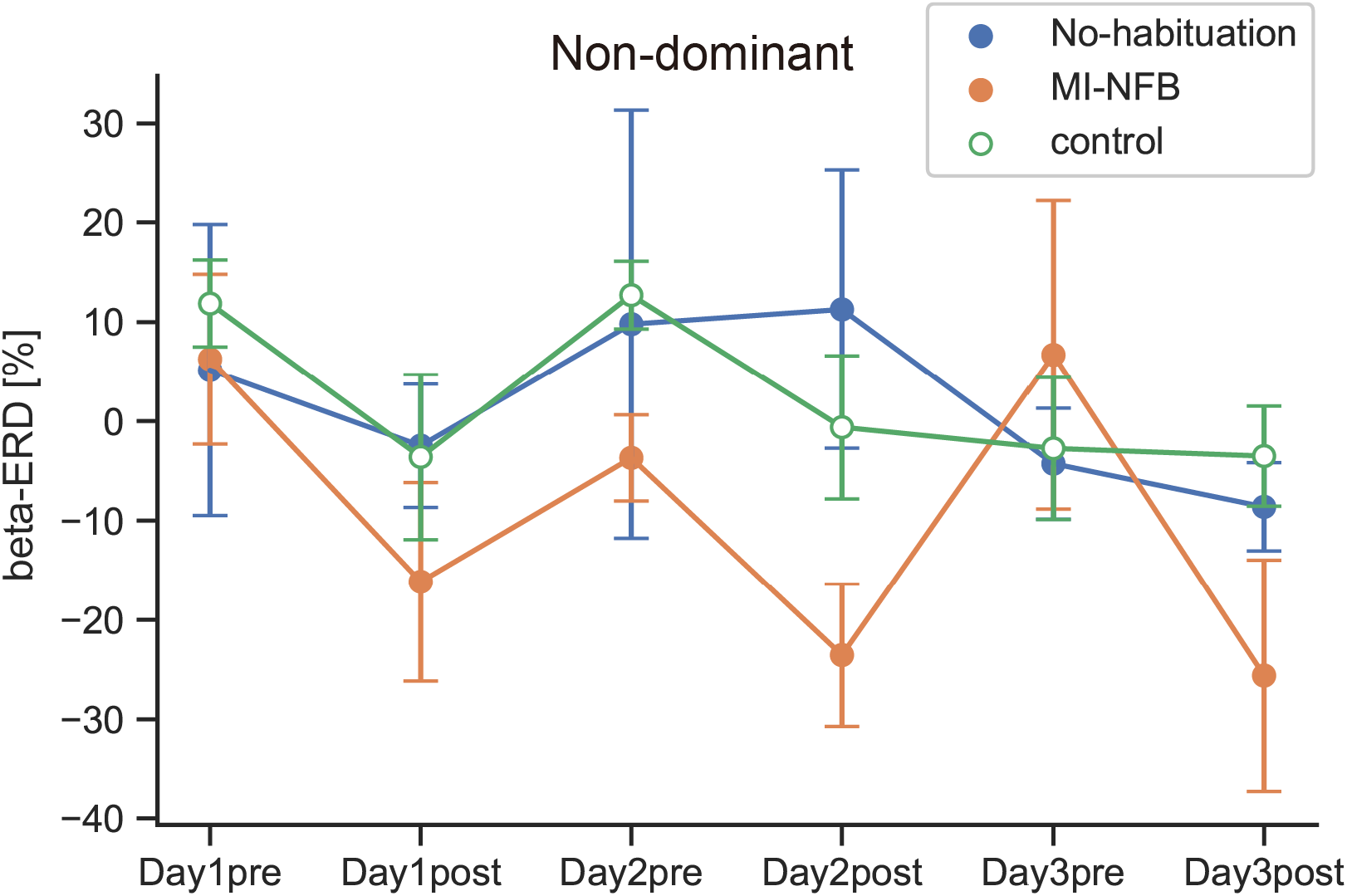
Results of beta-ERD with the non-dominant hand Mean and SEM in the mu-ERD from the pre-training session on day 1 to post-training on day 3. ERD: event-related desynchronization, MI-NFB: mental imagery-neurofeedback

### MI vividness

To evaluate the effect of MI-NFB on the MI subjective scale, the confidence rates were measured. The self-confidence rate of the MI-NFB group was significantly lower than that of the control group (*P* = 0.012).

To evaluate the effect of MI-NFB on MI ability and KVIQ scores were analyzed after the training. These measurements showed no significant differences between the MI-NFB and control groups, including No-habituation participants at post-measurement, VIS (t = −0.59, *P* =0.57), KIS (t = −0.88, *P* = 0.40), and SS (t = −0.77, *P* = 0.46).

### Experiment 2: Effects of MI-NFB on elite tennis athletes

In the second experiment, we investigated the effect of the proposed training on elite athletes. Five world-class male tennis players (T01–T05) were enrolled (Table 1). Because of the small number of target participants, we analyzed the training effects individually and did not perform any statistical analyses.

**Table 1.**
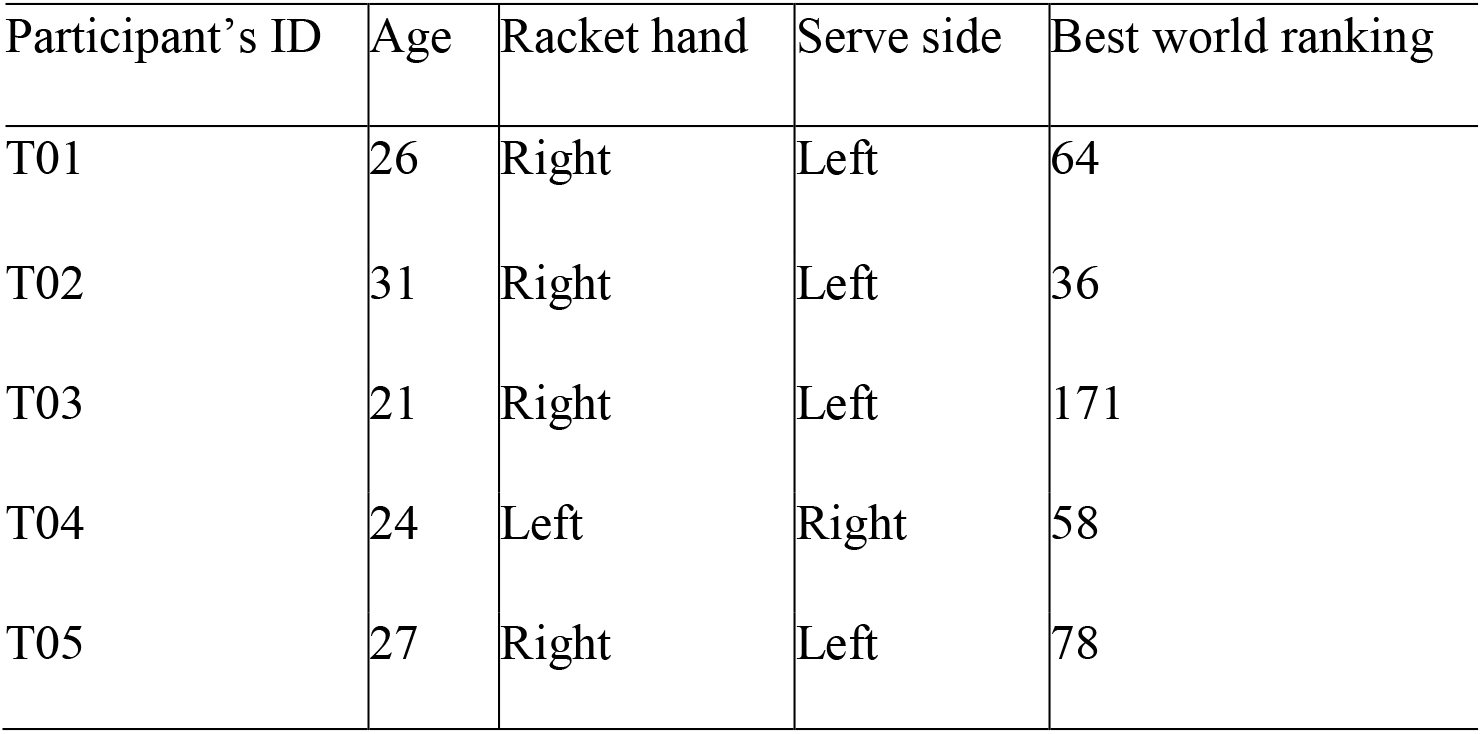
Demographic information of the tennis players.

| Participant's ID | Age | Racket hand | Serve side | Best world ranking |
| --- | --- | --- | --- | --- |
| T01 | 26 | Right | Left | 64 |
| T02 | 31 | Right | Left | 36 |
| T03 | 21 | Right | Left | 171 |
| T04 | 24 | Left | Right | 58 |
| T05 | 27 | Right | Left | 78 |

### Motor performance test

Motor performance was evaluated individually based on the number of serves that hit the target and the estimated service speed. Despite the lack of improvement in service speeds (Fig. 7B), the number of hits of three players increased from pre- to post-MI-NFB training. The number of hits of player T02 did not change, while the number of hits of player T03 decreased. (Fig. 7A). Motor performance was also evaluated subjectively based on the self-evaluation of each player for each service by the visual analog scale method. The subjective evaluations for services displayed an increase in players T02 and T04, while there was a decrease in players T03 and T05, and no changes in player T01 (Fig. 7C).

**Fig. 7.**
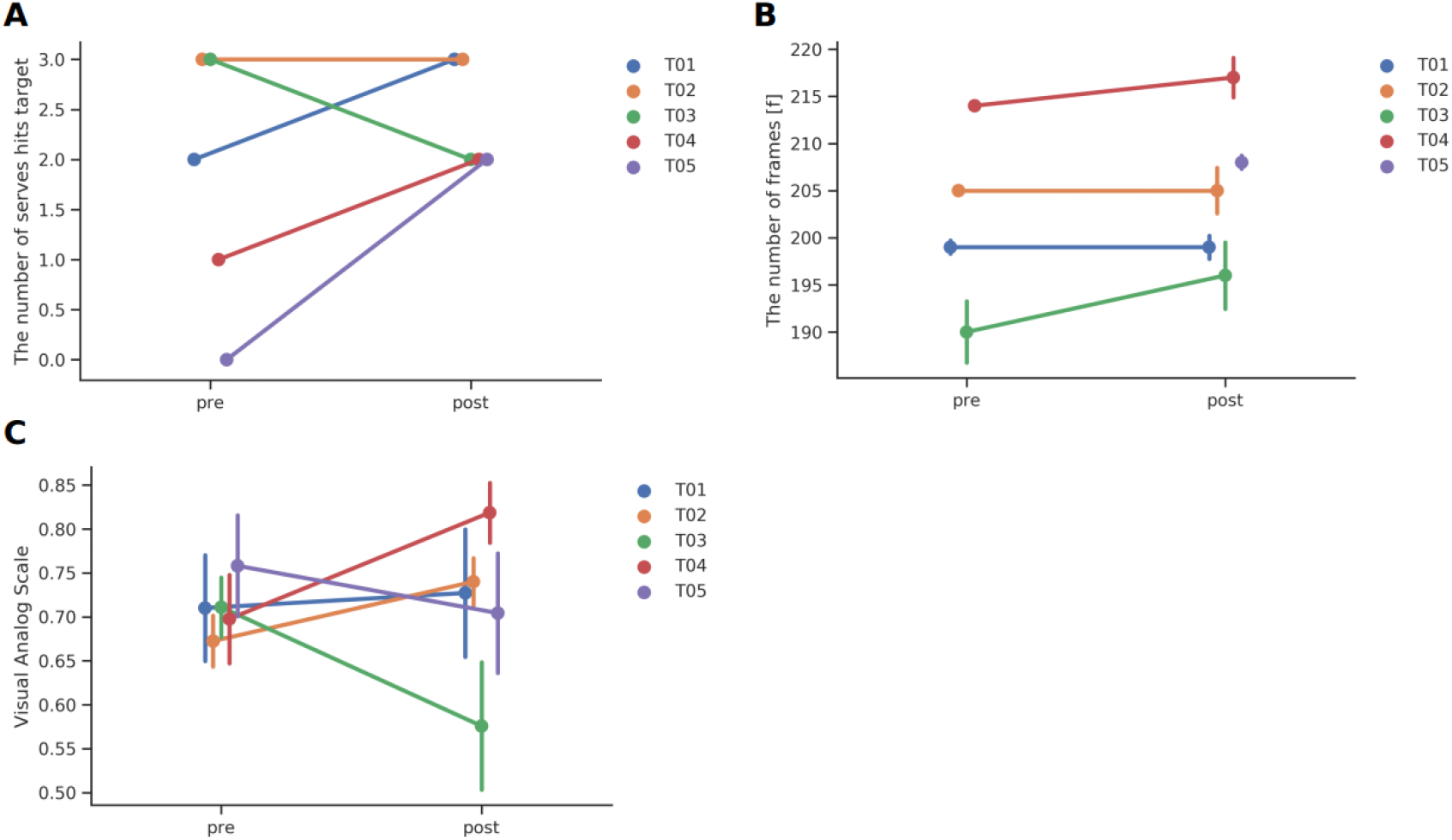
Results of experiments involving tennis athletes A: Changes in the number of hits on target before and after the MI-NFB training. B: Changes in the estimated speeds of serves before and after the MI-NFB training. C: Changes in subjective evaluation before and after the MI-NFB training. MI-NFB: mental imagery-neurofeedback

### The mu- and beta-ERD

To evaluate the training effect, relative mu-ERD and beta-ERD were analyzed by subtraction of pre-training on day 1 from post-training on day 3 in all electrodes. The topography indicated a grand average from right-handed athletes except for T04. The topography of mu-ERD showed enhancement after the training, especially in the ipsilateral side, though beta-ERD showed less enhancement than mu-ERD (Fig. 8).

**Fig. 8.**
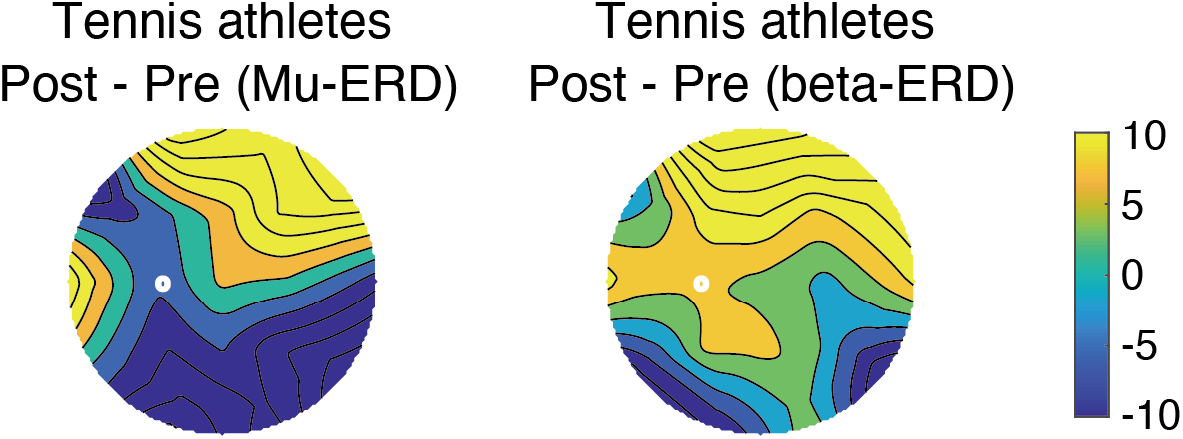
Topography of subtraction from post-training on day 3 to pre-training on day 1 The colors showed grand averaged mu-ERD (left) and beta-ERD (right) of right-handed tennis athletes. ERD: event-related desynchronization

## DISCUSSION

In this study, we aimed to enhance motor performance by using MI-based NFB training. We introduced an original performance test and developed a novel training protocol unifying all MI and NFB tasks into the original test. To verify the effectiveness of the proposed protocol, we compared motor performance following a training protocol with that following MI training alone. In Experiment 1, motor performance and beta-ERD amplitudes of participants who were trained under the protocol were more enhanced compared with those of the participants trained with MI alone, though we could not observe significant changes in mu-ERD. In Experiment 2, motor performances were improved in three of five athletes, and mu-ERD amplitudes of the athletes trained by the modified protocol for tennis players.

First, we discuss the different results between Experiment 1 and 2. Although both protocols at least partly enhanced motor performances, ERDs in different frequency bands were trained, mu or beta. This dissociated effectiveness in neuronal activity probably could be attributed to the participants’ familiarity levels of the motor action to learn. Both of mu- and beta-ERD are well known MI- or motor-related activities (Formaggio et al. 2010; Kraeutner et al. 2014; Nasseroleslami et al. 2014; Neuper et al. 2006; Gert Pfurtscheller and Neuper 1997; Sobierajewicz et al. 2017; Toriyama et al. 2018; Wheaton et al. 2009; Wolf et al. 2014; Wriessnegger et al. 2018). With regard to mu-ERD, larger mu-ERD has been observed more often in elite athletes than novices (Del Percio et al. 2007; Wolf et al. 2014). Beta-ERD has been discussed in the context of the motor learning and efficiency after learning (Gehringer et al. 2019; Jochumsen et al. 2017; Pollok et al. 2014). Beta-ERD has been related to the maintenance of a current motor state (Engel and Fries 2010), and, in one study, the short-term training was found to enhance beta-ERD, while longer-term training did not result in changes of beta-ERD compared with pre-training (Jochumsen et al. 2017). From these investigations, we can formulate the hypothesis that beta-ERD can be enhanced when learning an unfamiliar motor action, but it will be decreased after the action is familiarized. In line with this interpretation, a report has shown a correlation between the beta-ERD suppression and motor skill retention (Gehringer et al. 2018). In Experiment 1, we designed a novel motor task with short training periods. To acquire the new motor action, beta-ERD was modulated by the MI-related feedback. On the other hand, tennis services were very familiar among tennis elite athletes, and beta-ERD did not changed. The protocol used in Experiment 2 would enhance motor-related cortical activity by a different mechanism than that in Experiment 1, and it resulted in the enhancement of the motor performance and mu-ERD. The different frequency band of enhanced ERD might reflect the different methods of learning unfamiliar and familiar motor actions.

We hypothesized a mechanism for the proposed protocol, as described in Fig. 9C, before we performed Experiment 1. According to our hypothesis, the enhanced motor performance is derived from the mu-ERD feedback specified by the MI strategy of the given motor action. The trainee can enhance their mu-ERD from MI with feedback using own mu-ERD. This hypothesis (Fig. 9C) seems to be acceptable in the context of motor learning of familiar actions like Experiment 2. The mu-ERD is believed to reflect motor attention and multisensory integration, which are necessary to perform tasks more efficiently (G Pfurtscheller and Lopes Da Silva 1999; Wolf et al. 2014). From these reports, the effects of our training in Experiment 2 could be interpreted as the enhancement of the mu-ERD, which means that enhancing motor attention and integrating multisensory information results in a more efficient performance of an imagined motor action, and that mu-ERD enhancement may have affected the high-performance of a familiar action such as tennis service.

**Fig. 9.**
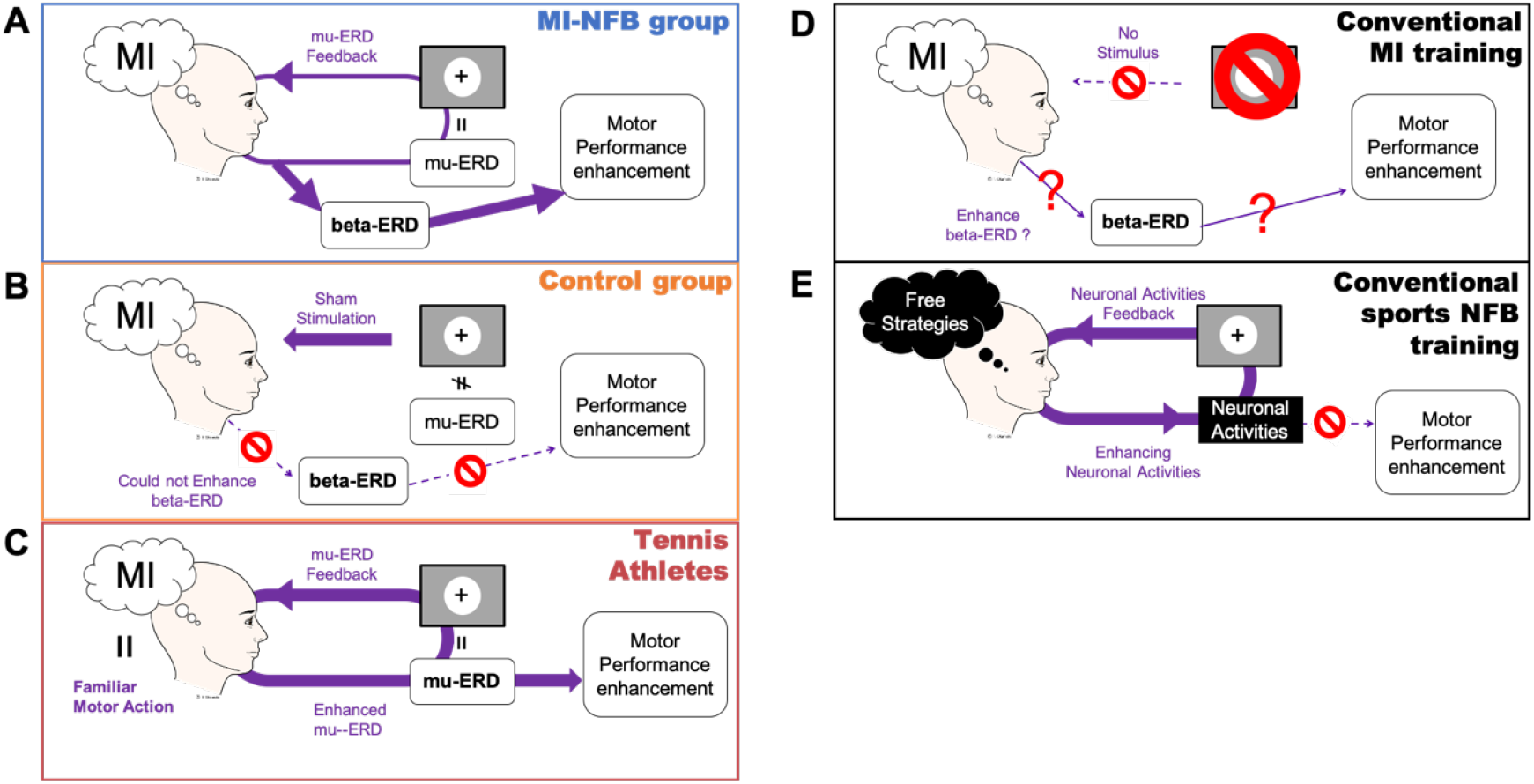
Conceptual scheme of the mechanism for the enhancement or non-enhancement of motor performance A: MI-NFB group with unfamiliar motor action. B: Control group. C: Tennis athletes with familiar motor action. D: Conventional MI training. E: Conventional NFB training. ERD: event-related desynchronization, MI: mental imagery, NFB: neurofeedback

From the results of Experiment 1, we hypothesized another mechanism for unfamiliar motor learning (Fig. 9A). The participants in MI-NFB were instructed to image the action as clearly as possible, and the feedback circle was presented as a marker of proficiency in the given motor action. Note that, they were not instructed to modulate the feedback activity itself. Thus, our results that beta-ERD enhancement resulted in the protocol using mu-ERD feedback would not contradict. The protocol used in Experiment 1 enabled enhancement of MI proficiency of a given action that was unfamiliar to the participants, and it provided enhancement of motor performance. We supposed that the beta-ERD enhancement resulted from the learning of an unfamiliar action.

Next, we discuss the group comparisons among the MI-NFB, no-habituation, and control groups, and subsequently, we interpret the results of the professional tennis athletes individually.

With regard to the MI-NFB and control groups in Experiment 1, we found that the performance of the dominant hand was enhanced after MI-NFB training. The participants could enhance their beta-ERD from MI using their own mu-ERD as the feedback. Several studies have reported that motor performance and ERDs of MI were positively correlated (Babiloni et al. 2008; Gibson et al. 2014). We can speculate that the mu-ERD feedback and strategy specification contributed to the success of the proposed training. In the control group, the motor performance was not enhanced, which suggests that participants were likely presented with feedback irrelevant to their own mu-ERD (Fig. 9B).

In conventional MI training studies, effects on motor performance are still controversial and may be attributable to the lack of mu-ERD feedback. The degree of MI vividness varied individually. Conventional studies did not involve any feedback of mu-ERD toward the degree of vividness, and the lack of the feedback might have provided ambiguous effect (Fig. 9D). In the conventional NFB-based studies on sports, motor performance could not be significantly enhanced, and beta-ERD was not trained by mu-ERD feedback. We speculate that there are two reasons for this: 1) the use of free strategies and 2) the irrelevant feedback to the motor action to be trained (Fig. 9E).

Contrary to our expectation, enhancement of motor performance while performing a motor action using the non-dominant hand was not observed after MI-NFB training. Generally, the efficiency of motor learning for a new action is lower in the non-dominant hand than in the dominant hand. We can expect that the participants failed to learn the unfamiliar action with the non-dominant hand using the training protocol. During the last evaluation session, no significant difference in working time was observed when an action was performed using the non-dominant hand. These results indicate that the training effect was not sustained until the last evaluation session. If we could prolong the training period long enough to sustain the beta-ERD enhancement until the last evaluation session, performance with the non-dominant hand would also likely be enhanced. Several reports have observed intermanual transfer, which is the ability to promote learning from a trained limb to another untrained limb using MI, and they observed the transfer effect from the trained non-dominant side to the untrained dominant side (Amemiya et al. 2010; Land et al. 2016; Van Mier and Petersen 2006). While these reports observed the transfer in keypad tapping or finger tracking, we employed an original peg-task including grasp, move, and insertion of small pegs. The complexity of a given action might have suppressed intermanual transfer from the dominant to the non-dominant hand. In the present experiment, we could not investigate the intermanual transfer effect precisely because both sides (dominant and non-dominant) underwent training. To analyze intermanual transfer in the proposed protocol, it is necessary to train only one side and examine the effect on the untrained side.

Self-confidence rates of MI were significantly higher in the control group than in the MI-NFB group. These results indicate that there was a gap between subjective evaluation and objective evaluation of MI. This cognitive gap was suggestive of a difficulty in estimating the ability of MI by subjective assessment alone.

KVIQ was conducted to investigate the effects on the abilities of general MI for any motor action by the training. There was no significant increase of KVIQ after the training. The result indicated that the general MI abilities were not affected by the training in the proposed protocol.

The participants in the no-habituation group who did not experience decreased mental fatigue during the training did not show enhanced motor performance and mu-ERD. Several previous studies have reported that high mental fatigue worsens motor performance. The mu-ERD of the no-habituation group was not enhanced by training. This result indicates that the four participants in the no-habituation group could not adapt to the NFB training. The stress of not being able to adapt to the NFB training caused mental fatigue, which may have worsened the motor performance. Therefore, controlling the mental fatigue of trainees may be necessary for successful NFB training.

Experiment 2 with tennis athletes was performed to assess the extendibility of the proposed training protocol. The training protocol was modified for tennis services. As a result, we found some training effects in most of the players. It is worth noting that our purpose was to enhance motor performance, not to improve MI vividness or MI-related ERD. In this experiment, three of five athletes could enhance their motor performance of services. Contrary to Experiment 1, mu-ERD was enhanced after the training, while beta-ERD was not. As we discussed above, we speculated that the mu-ERD enhancement reflected the proficiency of MI and performance of the familiar action.

We performed Experiment 2 as a case study, thus we discuss the results individually as follows. Player T04 showed remarkable effects which included: 1) the number of services that hit the target and 2) the values of subjective evaluation increased.

For players T01 and T05, the number of serves that hit the target increased after the training. In contrast, a decline in performance and ball speed was observed in player T03. Just before the post-performance test, player T03 could not do a warm-up exercise like the other participants; thus, the reduced warming up may have been the cause of the decline in performance. The service performance of player T02 did not differ from that of the pre-performance test. The mu-ERD of player T02 was not induced during training on day 3, likely because he could not train satisfactorily as a result of a system error. This failure to train mu-ERD on day 3 may have inhibited the enhancement of motor performance.

Our purpose in Experiment 2 was to investigate the extendibility of the proposed training protocol for elite athletes as a case report. This case report showed enhancement of motor performance in three athletes, and we suggest a possibility to use the protocol for field application, not only in laboratory experiments.

Wilson and Peper have hypothesized that elite athletes can benefit more from neurofeedback than non-athletes (Wilson and Peper 2011). Athletes are highly motivated to succeed in improvement and win. Feedback process is much easier for elite athletes because it is a familiar concept during training and practice. They can concentrate on many types of training with the belief that they will provide positive results. These characteristics and attitudes suggest the effectiveness of the proposed training.

It is useful to note that the protocol needs further modifications and investigations with a quantitative approach. More detailed measurements will provide a conclusion that the proposed training is effective for sports training.

The limitations of the study should be considered. In Experiment 1, the working time of the MI-NFB group decreased in the post-training more than that of the control group by mu-ERD feedback. These results strongly supported that the enhancement of motor performance was a result of MI-NFB training. However, we cannot reject the possibility that the motor performance was enhanced by the placebo effect and not by MI-NFB training itself. In our protocol of Experiment 1, we told the participants in the MI-NFB group that the visual stimuli were associated with their MI proficiency. Conversely, the participants of the control group were not told the meaning of the visual stimuli but were simply told to ignore them. This procedure was adopted to let the participants in the control group keep their motivation. Participants who received sham stimuli as feedback can easily notice their irrelevance, which lowers their motivation, as they were instructed to focus on the stimuli. However, the difference in instructions might have led to enhanced performance as a result of the placebo effect.

An additional limitation was the small sample size. In the training protocol (Experiment 1), the MI-NFB group was separated further by the cluster analysis due to their fatigue levels. This procedure was necessary to consider the relationship between training effect and habituation, but it inevitably reduced the sample size. Though the sample size was not sufficiently large, we observed some enhancement of motor performance in our protocol. Regarding the effects of MI-NFB on tennis athletes (Experiment 2), the results of the training in world-class tennis players indicated that the proposed training protocol could easily be applied to sports and could contribute to the enhancement of motor performance. We did not conduct group analysis in the tennis players experiment, because it was difficult to recruit enough world-class players. It is necessary to measure data from more elite players and to statistically compare the effect in a future study.

In both experiments, we did not measure electromyography, and we could not confirm whether the participants performed MI without the actual motion. We cannot reject the possibility that the observed mu-ERD was induced by an actual movement and not by imaging. We removed noisy data containing amplitudes exceeding 100 μV at Fp1. Most artifacts generated by body movements should have been rejected automatically, but it may not be enough to reject motor artifacts.

In conclusion, in the present study, we succeeded to enhance motor performance with the proposed training protocol using MI-NFB. We verified the hypothesis that the combination of MI and NFB would provide stable effects. The novelty of the proposed protocol was the specification of the strategy used for the MI of the intended motor action, which enabled feedback of EEG related to the MI and enhanced its performance. Notably, the protocol was effective for the trainees who did not experience increased mental fatigue during the training. If mental fatigue can be controlled appropriately, the proposed protocol can contribute to the development of more effective methods with applications in therapy and sports.

## MATERIALS AND METHODS

### Experiment 1: Training protocol for healthy volunteers

#### Participants

Eighteen Kyushu University students (9 women, 9 men; mean ± standard deviation (SD) aged 20.3 ± 1.7 years old; right-handed) were recruited for this study. One female participant was removed from our analysis because she could not perform our MPT due to an injury to her left thumb. Therefore, we analyzed the data from 17 participants (8 women; age, 20.4 ± 1.7 years old; right-handed). All participants performed the Flanders handedness test in Japanese to assess individual handedness (Okubo et al. 2014). All participants were healthy and had no previous history of neurologic disease or history of participating in similar MI experiments. All participants provided written informed consent. The experiments including any relevant details were approved by the local ethics committee of the Faculty of Arts and Science, Kyushu University. All experiments were performed in accordance with relevant guidelines and regulations.

Participants were divided into two groups, the MI-NFB group (5 women, 4 men; 20.3 ± 1.9 years old) and the control group (3 women, 5 men; 20.5 ± 1.6 years old), to avoid any biases of age (*t*(15) = −0.20, *P=*0.85), sports experience (*t*(15) = 0.68, *P =* 0.51), and the time period after retirement from sports (*t*(15) = 6.39, *P =* 0.70). One participant in control group was removed in EEG analysis because of the measurement problem on day 1.

#### Experimental training procedure

The participants sat on a comfortable armchair in an air-conditioned soundproof room. A monitor was placed at approximately 1.2 m in front of the participants (1920 × 1080 pixels).

The participants were asked to come to the experimental room for three days over a one-week period. On the first day, all participants engaged in a pre-test session, to undergo the assessment of motor performance; an evaluation session; a training session; and another evaluation session. On the second day, they participated in the first evaluation session, training session, and second evaluation session. On the third day, they first participated in the evaluation session, and then in the training session, second evaluation session, and post-test session for the assessment of motor performance.

#### Motor performance test (MPT)

We designed an MPT to control for unfamiliarity of the bilateral performance using a pegboard, similar to that used in previous studies for rehabilitation scenarios (Braun et al. 2017; Opara et al. 2017). In the performance test, the participants were instructed to insert four pegs into four different holes on the right or left side, as quickly as possible.

Immediately after the insertion, they were required to perform a 180° rotation of the pegs and to reinsert them in the holes located on the opposite side. Finally, they removed all pegs from the holes and pressed a key to measure the performance time (Fig. 10A). In the right-hand trial, in which the participants were directed to use their right hand, they moved the pegs from the right to the left side, and in the left-hand trial, they moved the pegs from the left to the right side. Each trial started with the presentation of an instruction to ‘Relax’ on the monitor for 5 s. An indication of ‘Left’ or ‘Right’ then appeared for 3 s to specify the lateral trial. Next, a fixation cross was presented, and the participants started the sequence of MPT. The fixation cross was presented while performing the task until a key was pressed. Before the beginning of the next trial, a blank screen re-appeared. The duration of the blank screen was randomly changed in the range of 2.1–2.5 s (Fig. 10C). Five trials of the performance test were conducted with each hand. The performance time was recorded from the time of presentation of the fixation cross to the moment of key pressing, and this performance time was used to evaluate motor performance bilaterally. A shorter performance time indicated high motor performance.

**Fig. 10.**
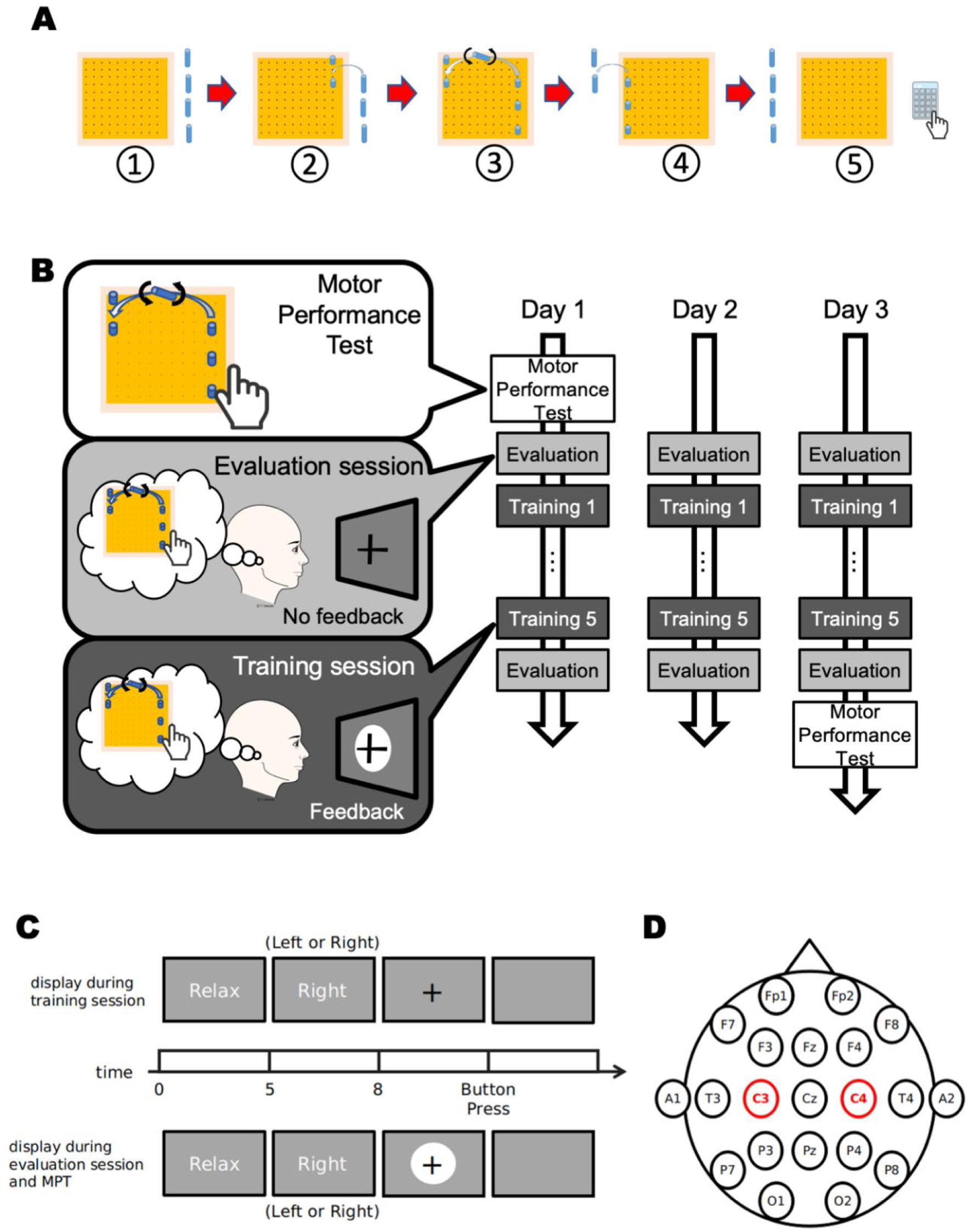
Experimental design of Experiment 1 A: Motor performance test (MPT) procedure. B: Procedure of the three training days. C: Procedure of each session. D: Electrode setup

#### Training session

In the MI-NFB training session, the participants performed MI using the same motor action as in the MPT. The displayed instructions and timings were designed to be the same as in the MPT trials (Fig. 10C). After laterality was specified, a circle was presented behind the fixation cross as a visual feedback, and the participants started MI. In the MI-NFB group, the radius of the circle varied according to their mu-ERD (see EEG analysis section below and equations 1 and 2). Participants were told that when they performed a better MI, the feedback circle would be smaller. They were instructed to attempt to make the circle smaller than the fixation cross, and as small as possible, by performing better MI. When the length of the fixation cross was same as the circle radius, it indicated 10% mu-ERD. In the control group, the radius of the feedback circle varied according to the mu-ERD of another participant used as sham stimulation. They were instructed to ignore the feedback circle and to concentrate on the MI.

A one-day training session consisted of five blocks, and one block included 5 MI trials for each hand (Fig. 10B). Therefore, a total of 50 MI trials were conducted in a training session. After each block, the participants took a 60-s break and answered a questionnaire using a 7-point scale indicating the difficulty of the MI (1: very easy, 7: very difficult), degree of mental fatigue (1: no fatigue, 7: high fatigue), and concentration (1: no concentration, 7: high concentration). Then they provided a self-confidence rate in the range of 0–100% for the performance of MI.

#### Evaluation session

In the evaluation session for the training effect, the participants performed MI as in the MPT trial but without any visual feedback (Fig. 10B). The display instructions and timing were the same as in the MPT trials (Fig. 10C). Participants in all groups were instructed to concentrate on MI by focusing on the fixation cross in each trial. There were no differences in instructions or display indications for all groups.

One evaluation session had a block including 5 MI trials for each hand, and this session was conducted before and after the one-day training session (Fig. 10B). After each evaluation session was completed, the participants took a 60-s break.

#### EEG recording

EEG data were recorded using Cognionics Quick-20 headsets (CGX, San Diego, CA, USA), consisting of 20 active-dry electrodes. These electrodes were placed according to the international 10-20 system, and the recorded signals were digitized with a sampling rate of 500 Hz. The reference electrode was placed on A1, at the left earlobe, and the ground electrodes were placed near Fp1 and Fp2 at the forehead. An additional electrode, A2 at the right earlobe was placed and measured (Fig. 10D). The active shield and active-dry electrodes in the Quick-20 enable active noise cancellation with high sensor impedances. The recommended sensor impedance of this device was below 2,500 kOhm, but it can tolerate up to 5,000 kOhm. In this study, impedance levels were maintained below 500 kOhm, which was 20% of the recommended level: 2500 kOhm. The impedance levels were monitored during the EEG recordings. We recorded EEG data during the training and evaluation sessions using a Data Acquisition Software (CGX, San Diego, CA, USA).

#### Clustering analysis according to performance and fatigue levels

Previous reports have pointed out that motor performance is impaired by mental fatigue (Marcora et al. 2009; Schiphof-Godart et al. 2018). To evaluate the training effect without impairment from mental fatigue, the participants who showed different fatigue patterns should be discriminated. To classify the participants, the fatigue levels and motor performance data were subjected to Ward’s hierarchical cluster analysis using Euclidean distances. The fatigue levels were recorded after each session every day, and we used the relative difference of fatigue data from day 1 to day 3 for clustering analysis, that is, five fatigue changes in every participant. The motor performance data were obtained by subtraction of working time on day 1 from day 3. The cluster analysis classified the data by fatigue changes and motor performances from day 1 to day 3.

#### EEG analysis

##### Real-time analysis

The decrease in real-time mu power from baseline (mu-ERD increased) was computed from the streaming EEG data. The recorded stream EEG data were stored for 1.024 s (512 points) before the onset of updating visual feedback, and the 1.024 s data were used to compute mu-ERD to define the radius of the feedback circle. We defined 8–13 Hz as the mu band. The baseline interval was defined as a relaxation period of 5 s just before a hand cue was presented (−8 s to −3 s before the hand cue onset), and the MI task interval was defined as the period from the presentation of the fixation cross to the moment of key pressing (0 to key pressing). We computed the mu-ERD using the following equation:

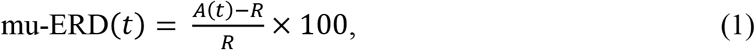

where *A*(*t*) is the power spectrum density of the time window during an MI trial at time *t*, and *R* is the baseline power at the trial. The radius of the feedback circle was computed from mu-ERD by using the following equation:

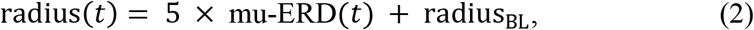

where radius_BL_ = 300 pixels. The visual angle when the radius_BL_ was presented was approximately 10.5°. The analytic time window during an MI trial was a sliding window with approximately 90% overlap. Accordingly, the feedback was updated approximately every 100 ms, and the analytic time window during the relaxation periods (baseline interval) was the sliding window without overlapping. The data in the baseline interval was computed as mu power in each window and averaged. The averaged data was used for the baseline power calculation (*R* in equation 1). The mu power spectrum density was computed by using the ‘psd_array_multitaper (array data, sampling rate)’ function in the Python MNE module. To remove eye-blinking artifacts, the time windows containing data with absolute values exceeding 100 μV at the Fp1 electrode were rejected. If the 1-s feedback was rejected because of the occurrence of artifacts during an MI trial, the data before the last feedback was used to compute the radius of the visual circle. Therefore, the size of the circle did not change when the data feedback contained artifacts. The data recorded at C3 were used to compute the mu-ERD of the right-hand MI, and the data at C4 were used to compute the mu-ERD of the left-hand MI. These analyses and feedback systems were developed using Python.

##### Offline analysis

Raw EEG data were band-pass filtered (3–50 Hz) and notch-filtered at 60 Hz. The recorded signals were re-referenced to the average of A1 and A2. The data were divided by −8 s to 15 s after the onset of MI start as one epoch. The mu-ERD was computed using equation (1). The EEG power from −6 s to −3 s was used as the baseline (*R*), and the power from 1 s to 5 s was used as an MI task activity (*A*). The remaining data from 5 s to 15 s after the MI onset was removed to exclude movement-related artifacts and EEG activity of the motor preparation for the button pressing.

#### Statistical analysis

In Experiment 1, we compared the data among the three groups, MI-NFB, control, and non-habituation. The motor performance were compared by a repeated measures ANOVA (pre/post all training days × group), and we used a t-test as post-hoc test. The mu-ERD and beta-ERD during the evaluation session was assessed by a repeated measures ANOVA (pre/post in each training day × group), and the t-test was used a post-hoc test. The significance α-value was set to 0.05, and Bonferroni correction was used for all statistical analyses. These tests were conducted in SciPy module (ver. 1.2.1) in Python (ver. 3.7.2).

### Experiment 2: Effects of MI-NFB on tennis athletes

#### Participants

Five world-class male tennis players were enrolled, and they ranked in the top 200 of Association of Tennis Professionals (ATP) men’s tennis ranking on May 13, 2019. The demographic data are summarized in Table 1.

Because of the inclusion of a small number of targeted participants, we analyzed the obtained data individually and did not perform a group comparison.

#### Experimental procedure

Only a few modifications were made to the experimental procedure used for healthy volunteers. The MPT was designed for tennis athletes as follows.

Training effects were evaluated by the performance of tennis serves. All participants hit 10 services before and after the training the right or left side on a tennis court. After each service, a participant evaluated the service subjectively using a visual analog scale ranging from “worst performance” to “best performance.” In the MPT, a service side was specified for each participant, from which he was not good at shooting a serve. The service side to shoot was determined according to the pre-training questionnaire of each participant. To evaluate the service performances, an 81 cm × 81 cm target area was designated on the tennis court (Fig. 11A). All players were instructed to hit the target area. The players’ motions and ball trajectories during services were recorded with three cameras from different locations. Ball speeds were estimated from videos recorded in 480 fps using the number of frames between the hit ball and grounded ball. We evaluated motor performance using the number of hits on target and estimated ball speeds. After all tasks were completed in the training protocol, we interviewed the participants.

**Fig. 11.**
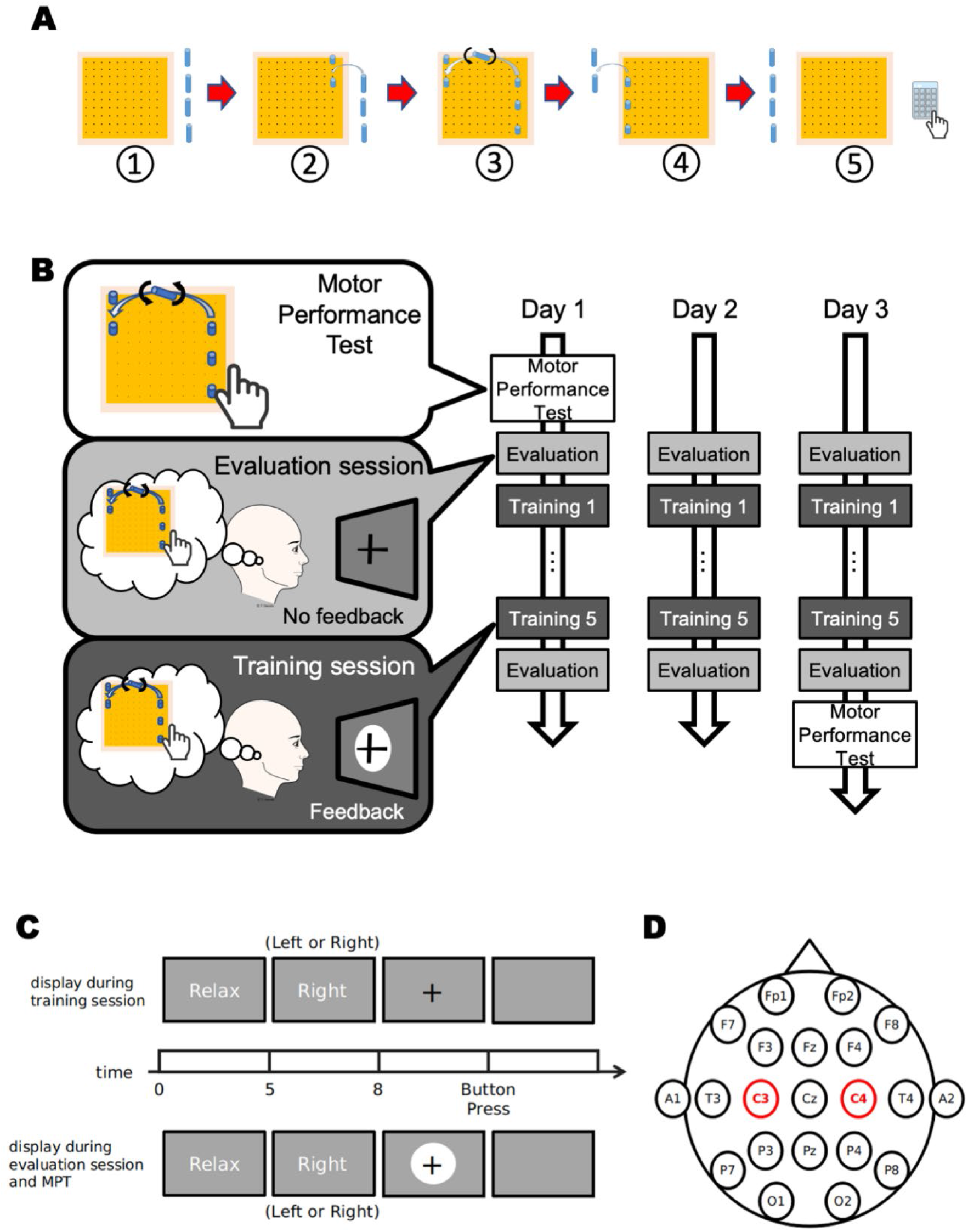
Experimental design of Experiment 2 A: Location of target of service, which was performed as a motor performance test (MPT). B: Procedure of the three training days

#### MI-NFB training

The MI-NFB training procedure was the same as that used for healthy volunteers. For the MI task in the athletes, participants were instructed to imagine the motion of serving from a side that was specified in the MPT (Fig. 11B). In this task, hand laterality of imaging was the same as the hand used by each participant to grip a racket.

#### Online analysis

On day 1 of the training protocol for athletes, the size of the feedback circle was computed by equation 1 as in the case of healthy volunteers, and the coefficient of ERD in equation 1 was changed to 4 on day 2 and to 3 on day 3. This change was intended to increase the difficulty of MI-NFB training, because we hypothesized that elite athletes have superior motor imagery ability.

## Acknowledgements

We thank Mr. K. Okuda (Universal Mind Inc.) and Mr. Y. Nishioka (Miki House Co., Ltd., professional tennis player) for their cooperation in scheduling and conducting the experiment on tennis athletes.

## Declarations

### Availability of data and material

The datasets generated during and/or analysed during the current study are available from the corresponding author on reasonable request.

### Code availability

None.

### Author Contributions

C.S. and T.O. conceived the idea and designed the study. C.S. and K.T. collected the data and performed the analysis. C.S. and K.T. wrote the draft with critical comments from T.O., and all authors discussed and finished the manuscript.

### Ethics approval

The experiments including any relevant details were approved by the local ethics committee of the Faculty of Arts and Science, Kyushu University (#201905, #20).

### Consent to participate

All participants provided written informed consent.

### Consent for publication

All authors consent for publication.

## References

Amemiya, K., Ishizu, T., Ayabe, T., & Kojima, S. (2010). Effects of motor imagery on intermanual transfer: A near-infrared spectroscopy and behavioural study. Brain Research, 1343, 93–103. https://doi.org/10.1016/j.brainres.2010.04.048

Arns, M., Kleinnijenhuis, M., Fallahpour, K., & Breteler, R. (2007). Golf performance enhancement and real-life neurofeedback training using personalized event-locked EEG profiles. Journal of Neurotherapy, 11(4), 11–18. https://doi.org/10.1080/10874200802149656

Babiloni, C., Del Percio, C., Iacoboni, M., Infarinato, F., Lizio, R., Marzano, N., et al. (2008). Golf putt outcomes are predicted by sensorimotor cerebral EEG rhythms. Journal of Physiology, 586(1), 131–139. https://doi.org/10.1113/jphysiol.2007.141630

Bai, O., Huang, D., Fei, D.-Y., & Kunz, R. (2014). Effect of real-time cortical feedback in motor imagery-based mental practice training. NeuroRehabilitation, 34(2), 355–363. https://doi.org/10.3233/NRE-131039

Boe, S., Gionfriddo, A., Kraeutner, S., Tremblay, A., Little, G., & Bardouille, T. (2014). Laterality of brain activity during motor imagery is modulated by the provision of source level neurofeedback. NeuroImage, 101, 159–167. https://doi.org/10.1016/j.neuroimage.2014.06.066

Braun, N., Kranczioch, C., Liepert, J., Dettmers, C., Zich, C., Büsching, I., & Debener, S. (2017). Motor Imagery Impairment in Postacute Stroke Patients. Neural Plasticity, 2017, 1–13. https://doi.org/10.1155/2017/4653256

Brown, T., Jamieson, G., & Cooper, N. (2012). Sensori-Motor Rhythm Neurofeedback Increases Fine Motor Skills in Elite Racket Sport Athletes. In ACNS-2012 Australasian Cognitive Neuroscience Conference. Frontiers Media SA. https://doi.org/10.3389/conf.fnhum.2012.208.00022

Caligiore, D., Mustile, M., Spalletta, G., & Baldassarre, G. (2017). Action observation and motor imagery for rehabilitation in Parkinson’s disease: A systematic review and an integrative hypothesis. Neuroscience and Biobehavioral Reviews, 72, 210–222. https://doi.org/10.1016/j.neubiorev.2016.11.005

Dal Maso, F., Desormeau, B., Boudrias, M. H., & Roig, M. (2018). Acute cardiovascular exercise promotes functional changes in cortico-motor networks during the early stages of motor memory consolidation. NeuroImage, 174, 380–392. https://doi.org/10.1016/j.neuroimage.2018.03.029

Decety, J., & Grèzes, J. (1999). Neural mechanisms subserving the perception of human actions. Trends in Cognitive Sciences, 3(5), 172–178. https://doi.org/10.1016/S1364-6613(99)01312-1

Del Percio, C., Brancucci, A., Bergami, F., Marzano, N., Fiore, A., Di Ciolo, E., et al. (2007). Cortical alpha rhythms are correlated with body sway during quiet open-eyes standing in athletes: A high-resolution EEG study. NeuroImage, 36(3), 822–829. https://doi.org/10.1016/j.neuroimage.2007.02.054

Di Rienzo, F., Debarnot, U., Daligault, S., Saruco, E., Delpuech, C., Doyon, J., et al. (2016, June 28). Online and offline performance gains following motor imagery practice: A comprehensive review of behavioral and neuroimaging studies. Frontiers in Human Neuroscience. Frontiers Media S. A. https://doi.org/10.3389/fnhum.2016.00315

Engel, A. K., & Fries, P. (2010). Beta-band oscillations — signalling the status quo? Current Opinion in Neurobiology, 20(2), 156–165. https://doi.org/10.1016/j.conb.2010.02.015

Formaggio, E., Storti, S. F., Cerini, R., Fiaschi, A., & Manganotti, P. (2010). Brain oscillatory activity during motor imagery in EEG-fMRI coregistration. Magnetic Resonance Imaging, 28(10), 1403–1412. https://doi.org/10.1016/j.mri.2010.06.030

Gehringer, J. E., Arpin, D. J., Heinrichs-Graham, E., Wilson, T. W., & Kurz, M. J. (2018). Neurophysiological changes in the visuomotor network after practicing a motor task. Journal of Neurophysiology, 120(1), 239–249. https://doi.org/10.1152/jn.00020.2018

Gehringer, J. E., Arpin, D. J., Heinrichs‐Graham, E., Wilson, T. W., & Kurz, M. J. (2019). Practice modulates motor‐related beta oscillations differently in adolescents and adults. The Journal of Physiology, 597(12), 3203–3216. https://doi.org/10.1113/JP277326

Gentili, R., Papaxanthis, C., & Pozzo, T. (2006). Improvement and generalization of arm motor performance through motor imagery practice. Neuroscience, 137(3), 761–772. https://doi.org/10.1016/j.neuroscience.2005.10.013

Gibson, R. M., Chennu, S., Owen, A. M., & Cruse, D. (2014). Complexity and familiarity enhance single-trial detectability of imagined movements with electroencephalography. Clinical Neurophysiology, 125(8), 1556–1567. https://doi.org/10.1016/j.clinph.2013.11.034

Gruzelier, J. H. (2014). EEG-neurofeedback for optimising performance. I: A review of cognitive and affective outcome in healthy participants. Neuroscience and Biobehavioral Reviews, 44, 124–141. https://doi.org/10.1016/j.neubiorev.2013.09.015

Hanslmayr, S., Sauseng, P., Doppelmayr, M., Schabus, M., & Klimesch, W. (2005). Increasing individual upper alpha power by neurofeedback improves cognitive performance in human subjects. Applied Psychophysiology Biofeedback, 30(1), 1–10. https://doi.org/10.1007/s10484-005-2169-8

Jeunet, C., Glize, B., McGonigal, A., Batail, J. M., & Micoulaud-Franchi, J. A. (2019). Using EEG-based brain computer interface and neurofeedback targeting sensorimotor rhythms to improve motor skills: Theoretical background, applications and prospects. Neurophysiologie Clinique, 49(2), 125–136. https://doi.org/10.1016/j.neucli.2018.10.068

Jochumsen, M., Rovsing, C., Rovsing, H., Cremoux, S., Signal, N., Allen, K., et al. (2017). Quantification of movement-related EEG correlates associated with motor training: A study on movement-related cortical potentials and sensorimotor rhythms. Frontiers in Human Neuroscience, 11, 604. https://doi.org/10.3389/fnhum.2017.00604

Kim, T., Frank, C., & Schack, T. (2017). A Systematic Investigation of the Effect of Action Observation Training and Motor Imagery Training on the Development of Mental Representation Structure and Skill Performance. Frontiers in Human Neuroscience, 11(October), 1–13. https://doi.org/10.3389/fnhum.2017.00499

Kober, S. E., Hinterleitner, V., Bauernfeind, G., Neuper, C., & Wood, G. (2018). Trainability of hemodynamic parameters: A near-infrared spectroscopy based neurofeedback study. Biological Psychology, 136, 168–180. https://doi.org/10.1016/j.biopsycho.2018.05.009

Kober, S. E., Pinter, D., Enzinger, C., Damulina, A., Duckstein, H., Fuchs, S., et al. (2019). Self-regulation of brain activity and its effect on cognitive function in patients with multiple sclerosis – First insights from an interventional study using neurofeedback. Clinical Neurophysiology, 130(11), 2124–2131. https://doi.org/10.1016/j.clinph.2019.08.025

Kober, S. E., Wood, G., Kurzmann, J., Friedrich, E. V. C., Stangl, M., Wippel, T., et al. (2014). Near-infrared spectroscopy based neurofeedback training increases specific motor imagery related cortical activation compared to sham feedback. Biological Psychology, 95(1), 21–30. https://doi.org/10.1016/j.biopsycho.2013.05.005

Kraeutner, S., Gionfriddo, A., Bardouille, T., & Boe, S. (2014). Motor imagery-based brain activity parallels that of motor execution: Evidence from magnetic source imaging of cortical oscillations. Brain Research, 1588, 81–91. https://doi.org/10.1016/j.brainres.2014.09.001

Land, W. M., Liu, B., Cordova, A., Fang, M., Huang, Y., & Yao, W. X. (2016). Effects of physical practice and imagery practice on bilateral transfer in learning a sequential tapping task. PLoS ONE, 11(4). https://doi.org/10.1371/journal.pone.0152228

Marcora, S. M., Staiano, W., & Manning, V. (2009). Mental fatigue impairs physical performance in humans. Journal of Applied Physiology, 106(3), 857–864. https://doi.org/10.1152/japplphysiol.91324.2008

Nan, W., Rodrigues, J. P., Ma, J., Qu, X., Wan, F., Mak, P. I., et al. (2012). Individual alpha neurofeedback training effect on short term memory. International Journal of Psychophysiology, 86(1), 83–87. https://doi.org/10.1016/j.ijpsycho.2012.07.182

Nasseroleslami, B., Lakany, H., & Conway, B. A. (2014). EEG signatures of arm isometric exertions in preparation, planning and execution. NeuroImage, 90, 1–14. https://doi.org/10.1016/j.neuroimage.2013.12.011

Neuper, C., Wörtz, M., & Pfurtscheller, G. (2006). Chapter 14 ERD/ERS patterns reflecting sensorimotor activation and deactivation. Progress in Brain Research. https://doi.org/10.1016/S0079-6123(06)59014-4

Okubo, M., Suzuki, H., & Nicholls, M. E. R. (2014). A Japanese version of the FLANDERS handedness questionnaire. Shinrigaku Kenkyu, 85(5), 474–481. https://doi.org/10.4992/jjpsy.85.13235

Opara, J. A., Małecki, A., Małecka, E., & Socha, T. (2017). Motor assessment in parkinson’s disease. Annals of Agricultural and Environmental Medicine, 24(3), 411–415. https://doi.org/10.5604/12321966.1232774

Paret, C., Goldway, N., Zich, C., Keynan, J. N., Hendler, T., Linden, D., & Cohen Kadosh, K. (2019). Current progress in real-time functional magnetic resonance-based neurofeedback: Methodological challenges and achievements. NeuroImage, 202(August), 116107. https://doi.org/10.1016/j.neuroimage.2019.116107

Park, J. L., Fairweather, M. M., & Donaldson, D. I. (2015). Making the case for mobile cognition: EEG and sports performance. Neuroscience and Biobehavioral Reviews, 52, 117–130. https://doi.org/10.1016/j.neubiorev.2015.02.014

Pfurtscheller, G, & Lopes Da Silva, F. H. (1999). Event-related EEG/MEG synchronization and desynchronization: Basic principles. Clinical Neurophysiology, 110(11), 1842–1857. https://doi.org/10.1016/S1388-2457(99)00141-8

Pfurtscheller, Gert, & Neuper, C. (1997). Motor imagery activates primary sensorimotor area in humans. Neuroscience Letters, 239(2-3), 65–68. https://doi.org/10.1016/S0304-3940(97)00889-6

Pollok, B., Latz, D., Krause, V., Butz, M., & Schnitzler, A. (2014). Changes of motor-cortical oscillations associated with motor learning. Neuroscience, 275, 47–53. https://doi.org/10.1016/j.neuroscience.2014.06.008

Rostami, R., Sadeghi, H., Karami, K. A., Abadi, M. N., & Salamati, P. (2012). The Effects of Neurofeedback on the Improvement of Rifle Shooters’ Performance. Journal of Neurotherapy, 16(4), 264–269. https://doi.org/10.1080/10874208.2012.730388

Ruffino, C., Bourrelier, J., Papaxanthis, C., Mourey, F., & Lebon, F. (2019). The use of motor imagery training to retain the performance improvement following physical practice in the elderly. Experimental Brain Research, 237(6), 1375–1382. https://doi.org/10.1007/s00221-019-05514-1

Ruffino, C., Papaxanthis, C., & Lebon, F. (2017). The influence of imagery capacity in motor performance improvement. Experimental Brain Research, 235(10), 3049–3057. https://doi.org/10.1007/s00221-017-5039-8

Schiphof-Godart, L., Roelands, B., & Hettinga, F. J. (2018). Drive in sports: How mental fatigue affects endurance performance. Frontiers in Psychology, 9(AUG), 1–7. https://doi.org/10.3389/fpsyg.2018.01383

Schuster, C., Hilfiker, R., Amft, O., Scheidhauer, A., Andrews, B., Butler, J., et al. (2011). Best practice for motor imagery: A systematic literature review on motor imagery training elements in five different disciplines. BMC Medicine, 9, 75. https://doi.org/10.1186/1741-7015-9-75

Sobierajewicz, J., Przekoracka-Krawczyk, A., Jaśkowski, W., Verwey, W. B., & van der Lubbe, R. (2017). The influence of motor imagery on the learning of a fine hand motor skill. Experimental Brain Research, 235(1), 305–320. https://doi.org/10.1007/s00221-016-4794-2

Spychala, N., Debener, S., Bongartz, E., Müller, H. H. O., Thorne, J. D., Philipsen, A., & Braun, N. (2020). Exploring Self-Paced Embodiable Neurofeedback for Post-stroke Motor Rehabilitation. Frontiers in Human Neuroscience, 13, 461. https://doi.org/10.3389/fnhum.2019.00461

Thompson, T., Steffert, T., Ros, T., Leach, J., & Gruzelier, J. (2008). EEG applications for sport and performance. Methods, 45(4), 279–288. https://doi.org/10.1016/j.ymeth.2008.07.006

Toriyama, H., Ushiba, J., & Ushiyama, J. (2018). Subjective vividness of kinesthetic motor imagery is associated with the similarity in magnitude of sensorimotor event-related desynchronization between motor execution and motor imagery. Frontiers in Human Neuroscience, 12(July), 1–10. https://doi.org/10.3389/fnhum.2018.00295

Van Mier, H. I., & Petersen, S. E. (2006). Intermanual transfer effects in sequential tactuomotor learning: Evidence for effector independent coding. Neuropsychologia, 44, 939–949. https://doi.org/10.1016/j.neuropsychologia.2005.08.010

Wheaton, L., Fridman, E., Bohlhalter, S., Vorbach, S., & Hallett, M. (2009). Left parietal activation related to planning, executing and suppressing praxis hand movements. Clinical Neurophysiology, 120(5), 980–986. https://doi.org/10.1016/j.clinph.2009.02.161

Wilson, V., & Peper, E. (2011). Athletes Are Different: Factors That Differentiate Biofeedback/Neurofeedback for Sport Versus Clinical Practice. Biofeedback, 39(1), 27–30. https://doi.org/10.5298/1081-5937-39.1.01

Wolf, S., BrÃ¶lz, E., Scholz, D., Ramos-Murguialday, A., Keune, P. M., Hautzinger, M., et al. (2014). Winning the game: brain processes in expert, young elite and amateur table tennis players. Frontiers in Behavioral Neuroscience, 8(October), 1–12. https://doi.org/10.3389/fnbeh.2014.00370

Wriessnegger, S. C., Brunner, C., & Müller-Putz, G. R. (2018). Frequency specific cortical dynamics during motor imagery are influenced by prior physical activity. Frontiers in Psychology, 9(OCT), 1–16. https://doi.org/10.3389/fpsyg.2018.01976

Zich, C., De Vos, M., Kranczioch, C., & Debener, S. (2015). Wireless EEG with individualized channel layout enables efficient motor imagery training. Clinical Neurophysiology, 126(4), 698–710. https://doi.org/10.1016/j.clinph.2014.07.007

Zich, C., Debener, S., Kranczioch, C., Bleichner, M. G., Gutberlet, I., & De Vos, M. (2015). Real-time EEG feedback during simultaneous EEG-fMRI identifies the cortical signature of motor imagery. NeuroImage, 114, 438–447. https://doi.org/10.1016/j.neuroimage.2015.04.020

Zich, C., Debener, S., Schweinitz, C., Sterr, A., Meekes, J., & Kranczioch, C. (2017). High-Intensity Chronic Stroke Motor Imagery Neurofeedback Training at Home: Three Case Reports. Clinical EEG and Neuroscience, 48(6), 403–412. https://doi.org/10.1177/1550059417717398

Zimmermann-Schlatter, A., Schuster, C., Puhan, M. A., Siekierka, E., & Steurer, J. (2008). Efficacy of motor imagery in post-stroke rehabilitation: A systematic review. Journal of NeuroEngineering and Rehabilitation, 5. https://doi.org/10.1186/1743-0003-5-8

