## Supplementary Figures for "Enhancement of motor performance by a training method combining motor imagery and neurofeedback"

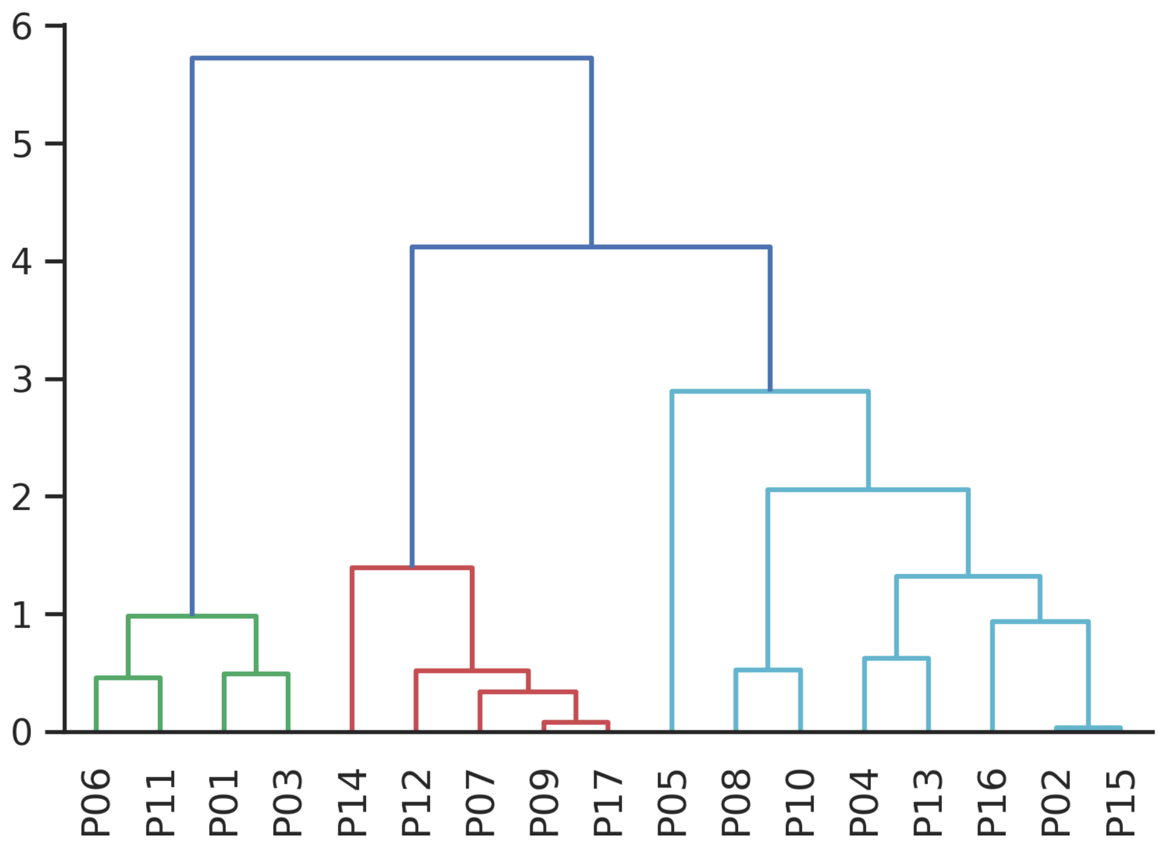


**Supplementary Figure 1. The dendrogram of a hierarchical clustering analysis of mental fatigue and MPT time.** From the analysis, P06, P11, P01, and P03, shown by the left green lines, were removed as outliers. They correspond to the four dots in the upper right of Fig. 5.2. MPT: motor performance test.

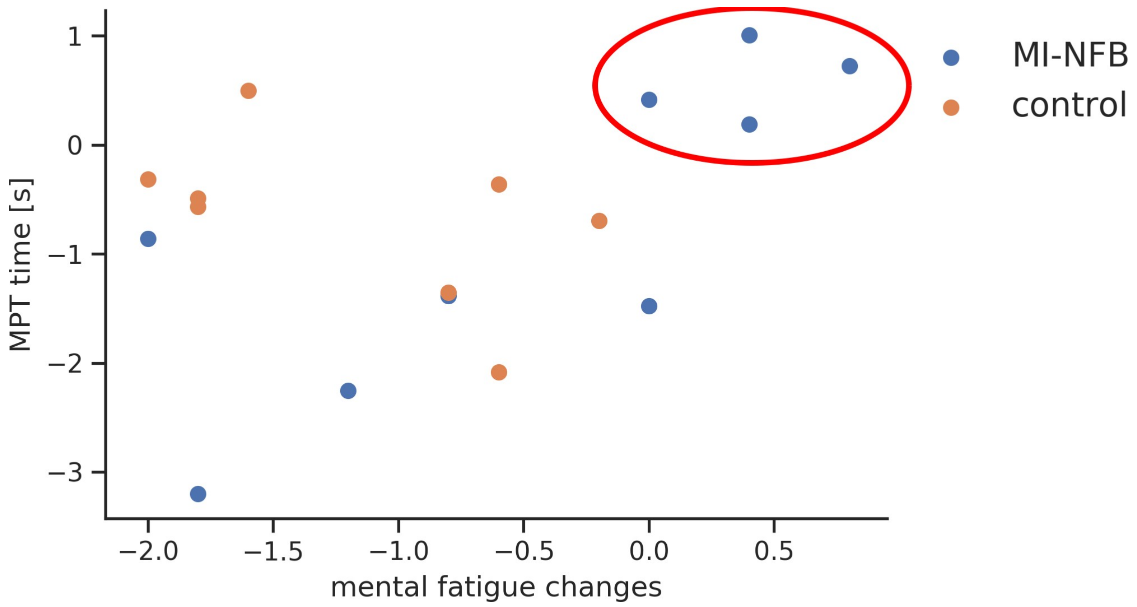


**Supplementary Figure 2. Scatter plots of mental fatigue changed according to MPT time. MPT: motor performance test, MI-NFB: mental imagery neurofeedback.**
